# A vast resource of allelic expression data spanning human tissues

**DOI:** 10.1101/792911

**Authors:** Stephane E. Castel, François Aguet, Pejman Mohammadi, GTEx Consortium, Kristin G. Ardlie, Tuuli Lappalainen

## Abstract

Allele specific expression (ASE) analysis robustly measures *cis* regulatory effects. Here, we present a vast ASE resource generated from the GTEx v8 release, containing 15,253 samples spanning 54 human tissues for a total of 431 million measurements of ASE at the SNP-level and 153 million measurements at the haplotype-level. In addition, we developed an extension of our tool phASER that allows effect sizes of *cis* regulatory variants to be estimated using haplotype-level ASE data. This ASE resource is the largest to date and we are able to make haplotype-level data publicly available. We anticipate that the availability of this resource will enable future studies of regulatory variation across human tissues.

## Background

Allele specific expression (ASE), or allelic expression analysis is a powerful technique that can be used to measure the expression of gene alleles relative to one another within single individuals. This makes it well suited to measure *cis*-acting regulatory variation using imbalance between alleles in heterozygous individuals (Fig. 1a) (Mohammadi, 2017). ASE analysis can capture both common *cis*-regulatory variation, for example, expression quantitative trait loci (eQTL), and rare regulatory variation (GTEx Consortium, 2017). It can also be used to measure allele-specific epigenetic effects such as parent of origin imprinting (Baran, 2015).

**Figure 1.**
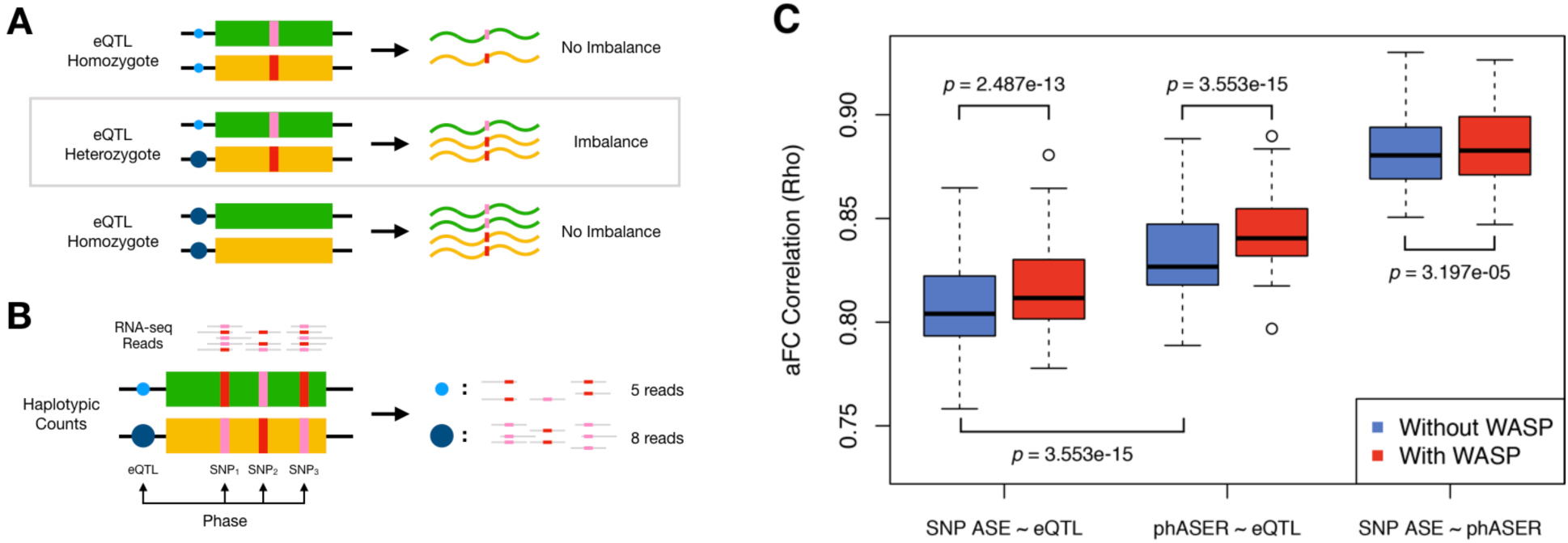
Capturing cis-regulatory effects with phased ASE data. **A)** The presence of a heterozygous *cis* regulatory variant or eQTL produces an expression level imbalance between the two haplotypes, which can be detected using allele-specific expression analysis. **B)** RNA-seq reads overlapping heterozygous SNPs in expressed regions of the gene can be used to quantify the expression of alleles relative to one another. These SNPs can be phased with each other and their counts aggregated to produce haplotype-level expression estimates, or haplotypic counts. The effects of regulatory variants can be captured by phasing them with haplotypic counts. **C)** Spearman correlation across 49 GTEx v8 tissues between eQTL effect size (allelic fold change, aFC) and effect size measured using ASE data from the single SNP with the highest coverage (SNP ASE) or haplotype-level ASE generated with phASER (phASER). Results are shown with and without allelic mapping bias correction from WASP. In each tissue, only a single top significant (FDR < 5%) eQTL per gene was analyzed. *p* values were calculated using a Wilcoxon paired signed rank test. For boxplots, bottom whisker: Q1 − 1.5*interquartile range (IQR), top whisker: Q3 + 1.5*IQR, box: IQR, and center: median.

In practice, ASE analysis uses RNA-seq reads that overlap heterozygous single nucleotide polymorphisms (SNPs), where the SNP can be used to assign the read to an allele. These heterozygous SNPs capture the cumulative effects of *cis*-regulatory variation acting on each allele. In some cases, these effects can be caused by the SNPs being used to measure ASE themselves, for example, stop gain variants that cause nonsense-mediated decay (NMD; Rivas, 2015), but often they simply capture effects of other *cis*-acting variation. Traditionally, a single SNP has been used to measure ASE, by taking the SNP with the highest coverage per gene. However, as a result of improvements in genome phasing, data can be aggregated across SNPs to produce estimates of allelic expression at the haplotype-level (Fig. 1b). We have previously developed a tool, phASER, which does this systematically, in a way that uses the information contained within reads to improve phasing, while preventing double counting of reads across SNPs to improve the quality of data generated (Castel, 2016).

In this work, we present an ASE resource generated using the Genotype Tissue Expression (GTEx) version 8 data release comprising RNA-seq data from 54 tissues and 838 individuals, for a total of 15,253 samples. We generated both SNP-level and haplotype-level ASE data. While the SNP-level data is available to approved users through dbGaP, the haplotype-level data does not contain identifiable information, and we were thus able to make it publicly available on the GTEx portal.

## Results and Discussion

Both SNP-level and haplotype-level ASE data were generated for each GTEx sample using current best practices, both with and without using WASP filtering (Geijn, 2015) to reduce the mapping bias that is sometimes present in ASE analysis, resulting in 4 data types per sample (Fig. S1, Methods – Data Generation). Across samples, this produced over 431 million measurements of ASE at the SNP-level and 153 million measurements of ASE at the haplotype-level. To demonstrate the ability of these data to robustly capture *cis*-regulatory effects and also benchmark the four data types relative to one another, we estimated eQTL effect sizes from ASE data using allelic fold change (aFC) (Mohammadi, 2017) and compared them to those derived from eQTL mapping (GTEx Consortium, 2019). To make it easier to generate aFC estimates for regulatory variants from phASER data, we developed a new add-on to the software package, phASER-POP, eliminating the need for custom scripts (Fig. S2, Methods – Software). We found high correlations between ASE and eQTL estimates, with a median spearman rho of 0.80 across tissues for SNP-level ASE data, and 0.83 for haplotype-level data generated by phASER (Fig. 1c). Haplotype-level correlations were significantly higher than SNP-level correlations (*p* = 3.55e-15, Wilcoxon paired signed rank test) while at the same time producing estimates for a median of 20% more genes (Fig. S3). WASP correction significantly improved correlations for both SNP-(*p* = 2.49e-13) and haplotype-(*p* = 3.55e-15) level data, so we therefore recommend using WASP-corrected data for most downstream analyses.

In the GTEx RNA-seq data, at a minimum coverage of 8 reads, samples had a median of 7,607 genes with ASE data at the SNP-level and 10,043 genes at the haplotype-level, and this dropped as a function of increasing coverage thresholds (Fig. S4). With the same coverage threshold, at the tissue level, there were a median of 18,042 genes with a median of 128 samples per gene using haplotype ASE data, rendering the data set well-powered to detect *cis*-regulatory effects (Fig. 2a). The median number of samples with ASE data per gene was largely dependent on tissue sample size, ranging from 39 for kidney (N = 73 samples) to 321 for thyroid (N = 574 samples). The number of genes with ASE data was determined by both sample size and gene expression diversity in tissues, with the two cell lines having the lowest number of genes with ASE data (LCLs = 15,804, fibroblasts = 16,526) and testis having the largest number of genes with ASE data (21,952) despite an intermediate sample size of 322.

**Figure 2.**
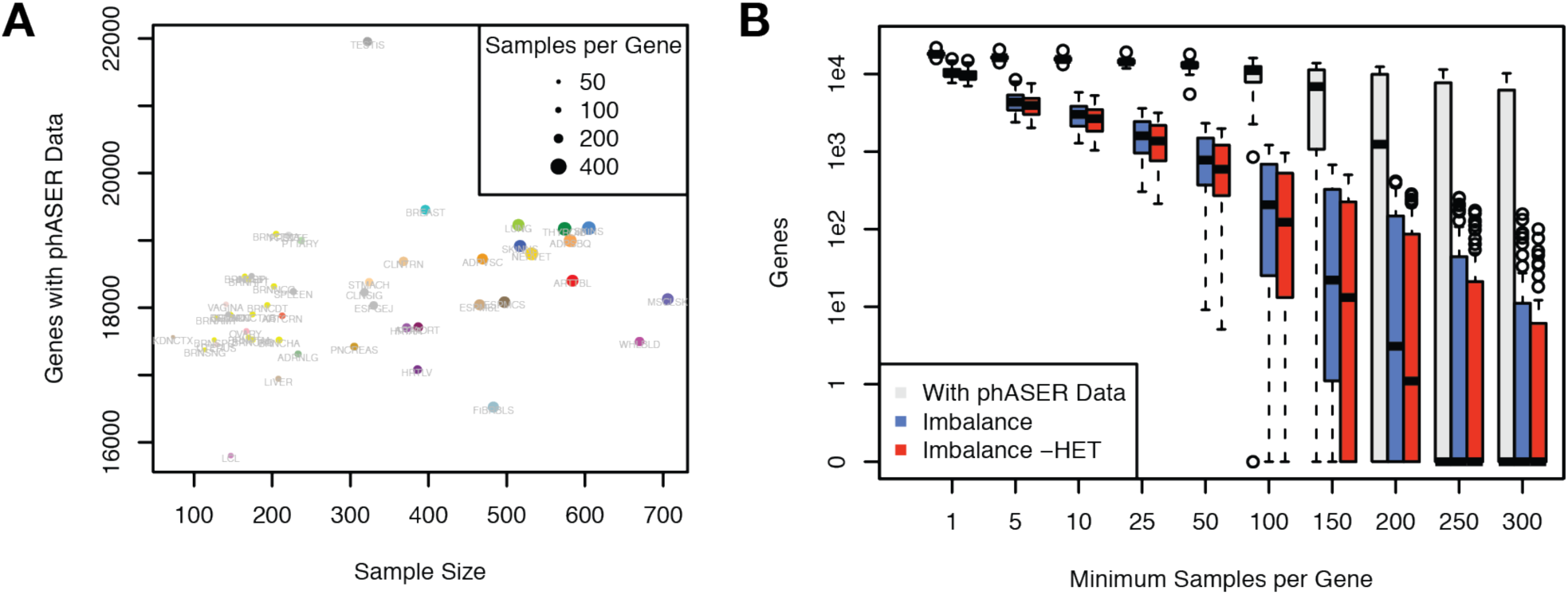
The GTEx v8 haplotype-level ASE resource. **A)** Number of genes per tissue with haplotype-level ASE data as a function of tissue sample size and the median number of samples with data per gene. **B)** Number of genes across the 49 GTEx tissues as a function of the minimum number of samples (*n*) with data per gene for: all genes with haplotype-level data in at least *n* individuals (grey), genes with significant (gene-level FDR < 5%) imbalance in at least *n* individuals (blue), genes with significant imbalance in at least *n* individuals who are not heterozygous for any top (FDR < 5%) or independent eQTL (permutation p < 1e-4). For boxplots, bottom whisker: Q1 − 1.5*interquartile range (IQR), top whisker: Q3 + 1.5*IQR, box: IQR, and center: median.

Finally, we sought to demonstrate the pervasiveness of *cis*-regulatory effects that can be captured with this resource. We found that even strong regulatory effects, where one allele was expressed at > 2x the level of the other allele, are widely present, even for protein coding genes, with 47% of protein coding genes showing such an effect in at least one tissue and at least 50 individuals (Fig S5). Considering all genes, we found that a median of 10,183 genes (or a median of 56% of those genes with ASE data) per tissue exhibited significant allelic imbalance (binomial test, FDR < 5% at the gene-level) in at least one sample, indicating the wide-spread nature of *cis*-regulatory effects (Fig. 2b). Removing individuals that were heterozygous for any known GTEx eQTL (Methods – GTEx eQTLs) only resulted in a median reduction of 7.5% in the number of genes with significant imbalance in at least one sample, demonstrating the potential of this resource to identify additional, likely rare regulatory effects that are not captured in eQTL analysis. A full summary of these statistics across tissues and sample thresholds is available in Table S1.

## Conclusion

In this work, we used the GTEx v8 release to produce a vast ASE resource, consisting of hundreds of millions of measurements of allelic expression. We generated SNP- and haplotype-level data, which provides better estimates of allelic expression for a greater number of genes. These data have numerous uses for the study of regulatory variation. For example, they have been used to replicate sex-, population- and cell type-specific eQTLs as well as capture the effects of rare regulatory variants (GTEx Consortium, 2019; Ferraro, 2019; Kim-Hellmuth, 2019). By making haplotype-level ASE data publicly available for the first time, we anticipate that this resource will find similarly broad use as the eQTL data it complements.

## Methods

### Data Generation and Availability

Paired-end 75bp Illumina RNA-seq reads were aligned to hg38 using STAR (Dobin, 2013) v2.5.3a (without allelic mapping bias correction) and v2.6.0c (with allelic mapping bias correction) in two-pass mode, and with allelic mapping bias correction enabled via the --waspOutputMode option which replicates the approach in van de Geijn *et al*., 2015 (the full settings of the alignment pipeline are described at https://github.com/broadinstitute/gtex-pipeline). All data was generated with or without using this feature and is indicated by “_WASP_” in the file names.

SNP-level ASE data was generated using the GATK ASEReadCounter tool v3.8-0-ge9d806836 with the following settings: -U ALLOW_N_CIGAR_READS -minDepth 1 -- allow_potentially_misencoded_quality_scores --minMappingQuality 255 --minBaseQuality 10. Raw SNP-level data, consisting of the GATK tool output, were aggregated per subject across all tissues. Raw autosomal SNP-level data, for SNPs with ≥8 reads, was annotated by: assigning heterozygous SNPs to genes using Gencode v26, calculating the expected null ratio for each combination of ref/alt allele (Castel, 2015), calculating a binomial p-value by comparing to the expected null ratio, calculating a multiple hypothesis corrected p-value per tissue using Benjamini-Hochberg, and flagging sites that overlapped low-mappability regions (75-mer mappability <1 based on 75mer alignments with up to two mismatches), showed mapping bias in simulation (Panousis, 2014), or had no more reads supporting two alleles than would be expected from sequencing noise alone, indicating potential genotyping errors (FDR < 1%, see Castel, 2015 for description of test). The genotype warning test cannot distinguish between strong allelic expression and a true genotyping error, and as a result should not be used when studying phenomena with expected mono-allelic expression (e.g. imprinting).

Haplotype-level data was generated using phASER v1.0.1 (Castel, 2016). phASER was run using whole genome sequencing genotype calls that were population-phased with Shapeit v2.837 in read-backed phasing mode with whole genome sequencing reads (Delaneau, 2013). phASER was run using all available RNA-seq libraries per subject. RNA-seq read-backed phased genotype data are provided (filename: phASER_GTEx_v8_merged.vcf.gz). Haplotypic expression was calculated using phASER Gene AE 1.2.0 and Gencode v26 gene annotations with min_haplo_maf 0.01. Haplotypic expression matrices containing all samples were generated using the “phaser_expr_matrix.py” script. This consists of a single string per sample per gene with the format “HAP_A_COUNT|HAP_B_COUNT”. One matrix was generated using only haplotypes that could be genome wide phased such that the haplotype assignment is consistent across genes within an individual and with the phased VCF (filename: phASER_GTEx_v8_matrix.gw_phased.txt.gz). Another was generated that does not ensure genome wide haplotype phasing across genes, which includes more counts, but makes the haplotype assignment of A/B arbitrary and unrelated across genes within an individual or the VCF (filename: phASER_GTEx_v8_matrix.txt.gz). The full settings of the haplotype-level ASE pipeline are described at https://github.com/broadinstitute/gtex-pipeline/.

SNP-level data is available for authorized users via dbGaP under accession phs000424 (filenames: phe000039.v1.GTEx_v8_ASE.expression-matrixfmt-ase.c1.GRU.tar, phe000039.v1.GTEx_v8_ASE_WASP.expression-matrixfmt-ase.c1.GRU.tar). phASER generated, haplotype-level data is available through the same dbGaP accession (folders GTEx_Analysis_v8_phASER and GTEx_Analysis_v8_phASER_WASP inside archive phe000037.v1.GTEx_v8_RNAseq.expression-data-matrixfmt.c1.GRU.tar) and on the GTEx Portal (http://gtexportal.org/).

### Software and Availability

The original phASER package produced gene-level haplotypic expression per individual (Castel, 2016). We developed new additions to phASER (phASER-POP) that make it easier to analyze data across many samples, as is often done with gene expression quantifications. First, we developed a new addition to the software (phaser_expr_matrix.py) that enables the aggregation of gene-level haplotypic expression measurement files across samples to produce a single haplotypic expression matrix, where each row is a gene and each column is a sample. The values consist of a single string per sample per gene in the format “HAP_A_COUNT|HAP_B_COUNT”. This format is intended to facilitate downstream analyses of allelic expression.

Second, we developed a tool to make it easier to estimate effect sizes of regulatory variants using phASER haplotypic expression data (phaser_cis_var.py). As input, this script takes a phASER haplotype expression matrix, a VCF, and a list of variants to calculate effect sizes for. To improve accuracy, the read-backed phased VCFs produced by phASER should be used, but first need to be combined across individuals, which can be performed using, e.g., “bcftools merge ind1.vcf.gz ind2.vcf.gz …”. The tool phases variant alleles with haplotype expression data and outputs numerous statistics, including allelic fold change (aFC, Mohammadi, 2017) per sample, and a median across samples for individuals that heterozygous for the variant of interest. This median can be used as the estimate of the regulatory variant effect size. The output also includes statistics calculated for homozygous individuals, which can be used to test for increased allelic imbalance in heterozygotes as compared to homozygotes, as would be expected for true regulatory variants.

The software along with extensive documentation is available through GitHub (http://www.github.com/secastel/phaser).

### GTEx eQTLs

For comparison between eQTL effect size and allelic expression effect size, GTEx v8 top significant (FDR < 5%) eQTLs were used from 49 tissues (GTEx Consortium, 2019). This results in at most a single eQTL per gene in a given tissue. When quantifying the number of samples that are not heterozygous for a known eQTL but still show allelic imbalance, gene-level haplotypic expression levels were excluded for a sample if the individual was heterozygous for a top significant eQTL or a nominally significant (permutation p < 1e-4) independent eQTL in any of the 49 tissues.

## Declarations

### Availability of data and materials

The datasets generated and/or analyzed during the current study are available to authorized users via dbGaP under accession phs000424 and on the GTEx portal (http://gtexportal.org/).

### Authors’ Contributions

S.E.C. and F.A. developed pipelines and generated data. S.E.C. and P.M. analyzed the data. S.E.C. developed the phASER software. S.E.C., K.A., and T.L. and designed the study. S.E.C., F.A., P.M., T.L., wrote and edited the manuscript.

## Acknowledgements

We would like to thank members of the Lappalainen laboratory for discussions surrounding the project. We thank the GTEx donors for their contributions to science, the GTEx Laboratory, Data Analysis, and Coordinating Center (LDACC), and the GTEx analysis working group (AWG) for their work in generating the resource.

## Supplemental Figures

**Figure S1.**
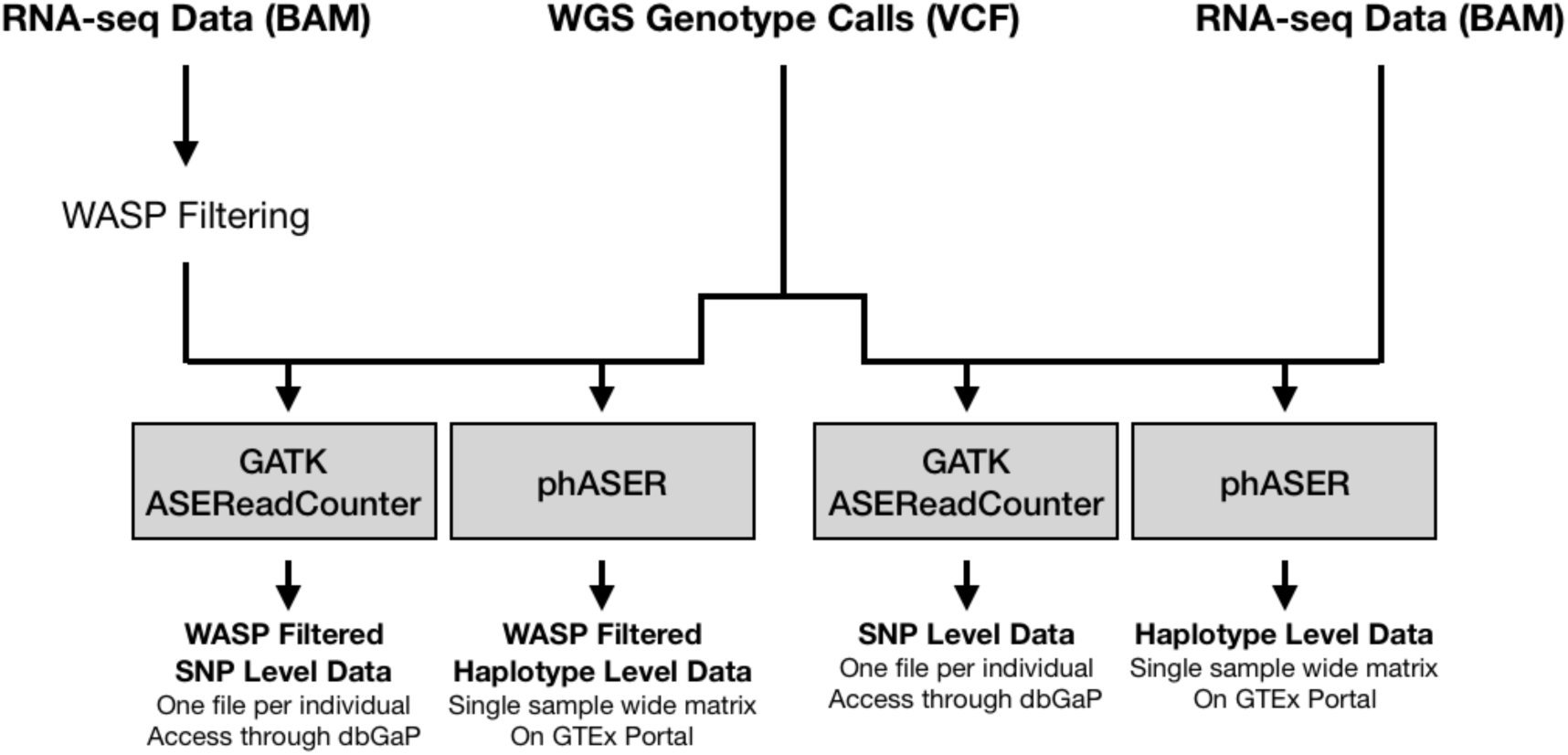
GTEx v8 ASE data types. ASE data is available at both the SNP and haplotype-level (single measurement per gene) with and without WASP filtering to reduce mapping bias. For SNP-level data, a single file was generated per individual containing data from across all tissues that were sampled from that individual and is available through dbGaP. At the haplotype-level, a matrix containing a single ASE measurement per gene across all GTEx samples is available publicly through the GTEx portal.

**Figure S2.**
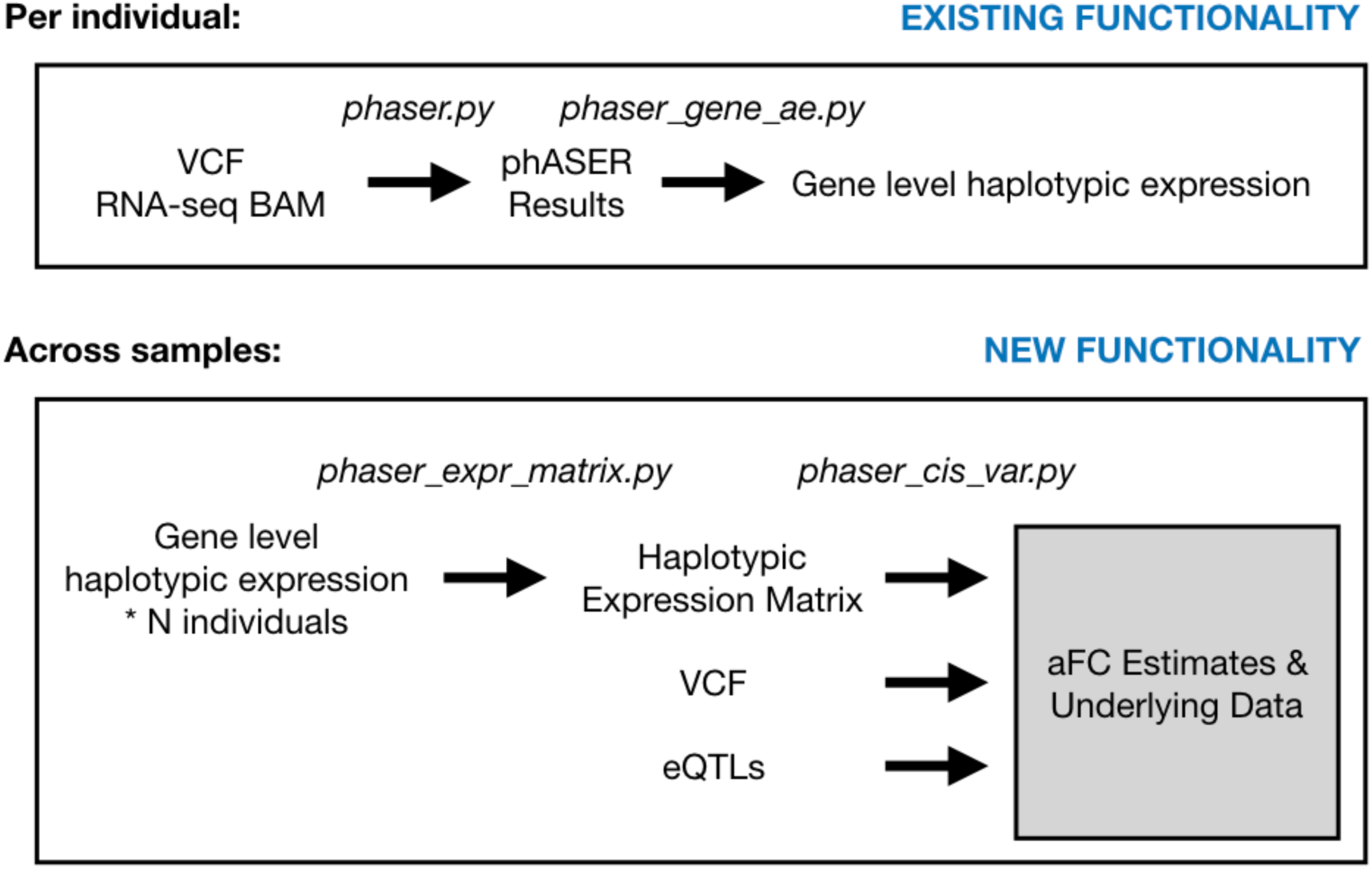
Additions to the phASER package that allow for easy measurement of regulatory variant effects using phased ASE data. The original phASER package produced gene-level haplotypic expression per individual. The new additions allow gene-level haplotypic expression measurement files to be combined across individuals to produce a single haplotypic expression matrix, where each row is a gene and each column is an individual. This matrix can then be used to retrieve ASE data and calculate relevant statistics for a given set of eQTLs or any other variants of interest.

**Figure S3.**
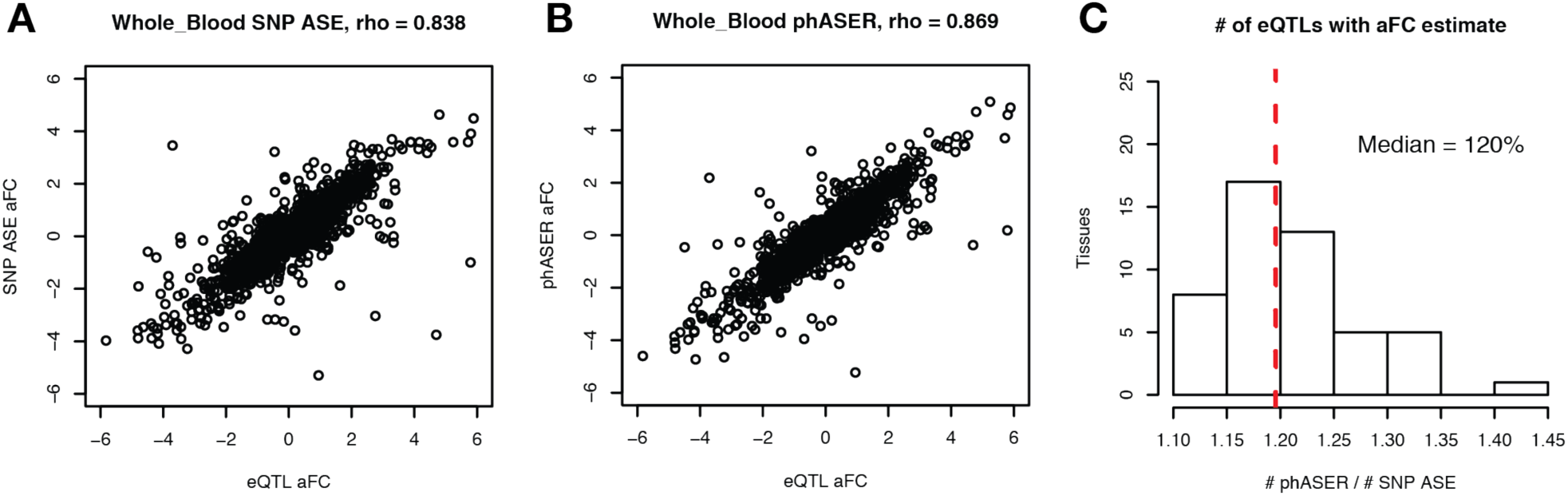
Capturing *cis*-regulatory effects with phased ASE data extended. **A-B)** Correlation between eQTL effect size (aFC) and effect size measured using ASE data from the single SNP with the highest coverage **(A)** or haplotype-level ASE generated with phASER **(B)** in GTEx v8 Whole Blood. **C)** Number of eQTLs with sufficient (≥ 10 individuals) allelic expression data to estimate effect size using phASER to generate haplotype-level estimates versus using the single SNP with the highest coverage.

**Figure S4.**
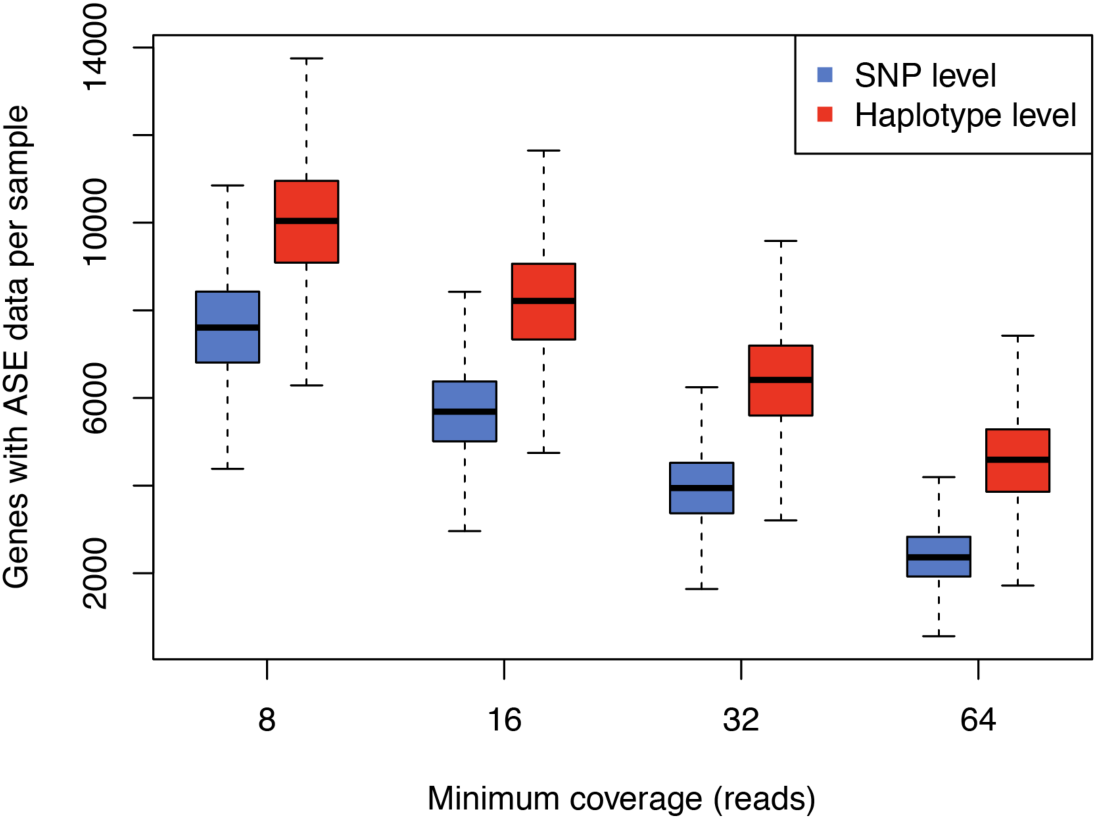
Sample level ASE data metrics. Boxplot showing number of genes with ASE data per sample across all GTEx tissues using either SNP-level (blue) or haplotype-level (red) data as a function of minimum read coverage. For boxplots, bottom whisker: Q1 − 1.5*interquartile range (IQR), top whisker: Q3 + 1.5*IQR, box: IQR, and center: median and outliers are hidden for ease of viewing.

**Figure S5.**
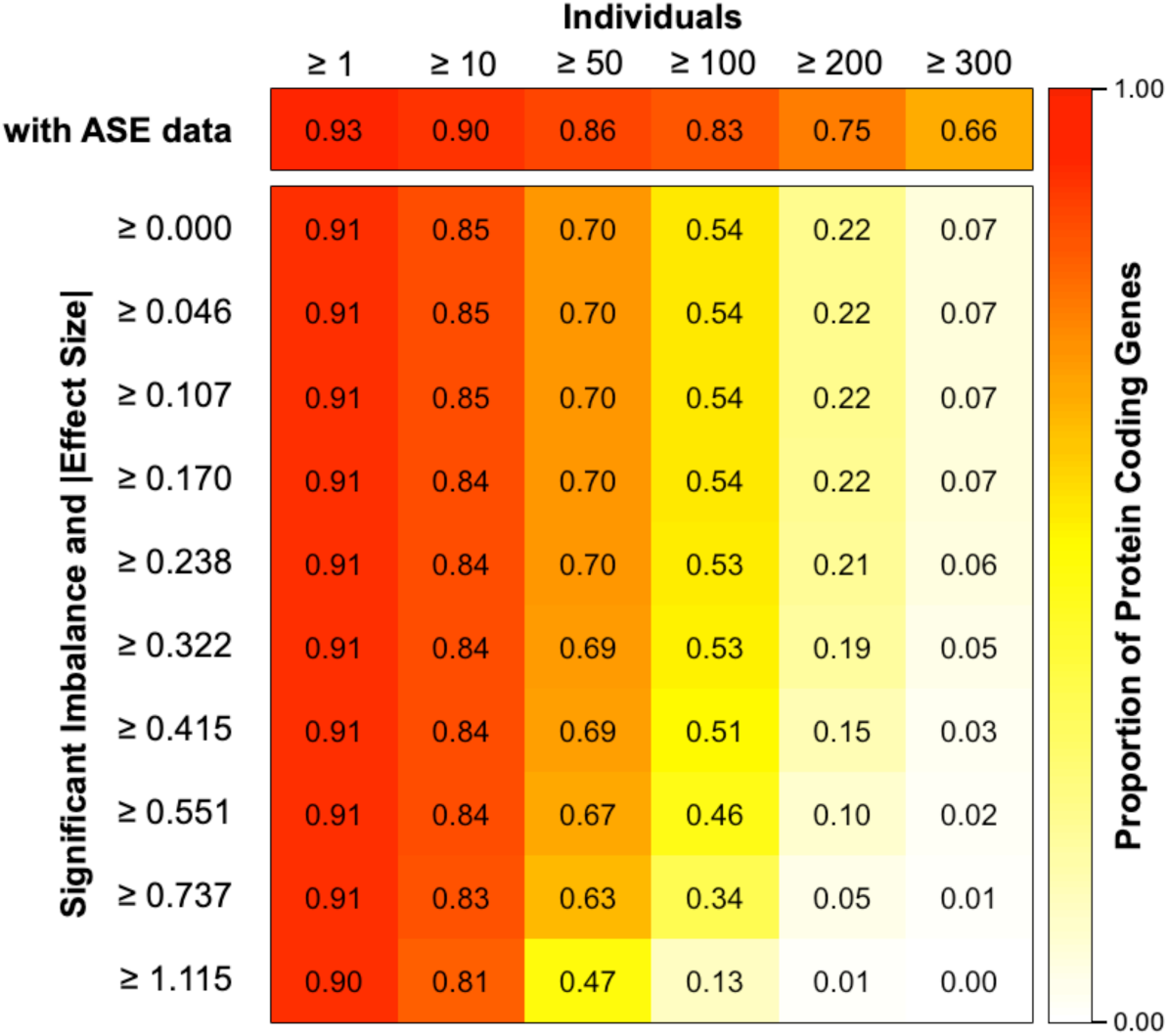
Phased ASE data at protein coding genes across tissues. Heatmap showing the proportion of protein coding genes with ASE data in at least one GTEx tissue as a function of the number of individuals with data using all ASE data (top row) or only those samples with significant imbalance (sample level FDR < 5%) and increasing minimum effect size, calculated using aFC.

## Supplemental Tables

**Table S1. GTEx haplotype-level ASE statistics**. Table where rows are each of the 49 GTEx tissues where eQTLs were called and columns list the number of genes with haplotype-level ASE data at minimum number of sample thresholds from 1 to 300 (minXXX). For example, min1 lists the number of genes that have ASE data from at least 1 sample. The table has three sheets, the first (all_data) presents statistics generated using all haplotype-level ASE data, the second (sig_imb_fdr05), counting only cases with significant allelic imbalance (binomial test versus 50/50, gene-level FDR < 5%), and finally (sig_imb_fdr05_no_het), counting only cases with significant imbalance where the individual is not heterozygous for any top (FDR < 5%) or independent (permutation p < 1e-4) eQTLs across any GTEx tissues for the gene.

